# Historical biogeography of the leopard (*Panthera pardus*) and its extinct Eurasian populations

**DOI:** 10.1101/413120

**Authors:** Johanna L.A. Paijmans, Axel Barlow, Daniel W. Förster, Kirstin Henneberger, Matthias Meyer, Birgit Nickel, Doris Nagel, Rasmus Worsøe Havmøller, Gennady F. Baryshnikov, Ulrich Joger, Wilfried Rosendahl, Michael Hofreiter

## Abstract

**Background:** Resolving the historical biogeography of the leopard *(Panthera pardus)* is a complex issue, because patterns inferred from fossils and from molecular data lack congruence. Fossil evidence supports an African origin, and suggests that leopards were already present in Eurasia during the Early Pleistocene. Analysis of DNA sequences however, suggests a more recent, Middle Pleistocene shared ancestry of Asian and African leopards. These contrasting patterns led researchers to propose a two-stage hypothesis of leopard dispersal out of Africa: an initial Early Pleistocene colonisation of Asia and a subsequent replacement by a second colonisation wave during the Middle Pleistocene. The status of Late Pleistocene European leopards within this scenario is unclear: were these populations remnants of the first dispersal, or do the last surviving European leopards share more recent ancestry with their African counterparts?

**Results:** In this study, we generate and analyse mitogenome sequences from historical samples that span the entire modern leopard distribution, as well as from Late Pleistocene remains. We find a deep bifurcation between African and Eurasian mitochondrial lineages (∼710 Ka), with the European ancient samples as sister to all Asian lineages (∼483 Ka). The modern and historical mainland Asian lineages share a relatively recent common ancestor (∼122 Ka), and we find one Javan sample nested within these.

**Conclusions:** The phylogenetic placement of the ancient European leopard as sister group to Asian leopards suggests that these populations originate from the same out-of-Africa dispersal which founded the Asian lineages. The coalescence time found for the mitochondrial lineages aligns well with the earliest undisputed fossils in Eurasia, and thus encourages a re-evaluation of the identification of the much older putative leopard fossils from the region. The relatively recent ancestry of all mainland Asian leopard lineages suggests that these populations underwent a severe population bottleneck during the Pleistocene. Finally, although only based on a single sample, the unexpected phylogenetic placement of the Javan leopard could be interpreted as evidence for exchange of mitochondrial lineages between Java and mainland Asia, calling for further investigation into the evolutionary history of this subspecies.

## Background

Achieving a comprehensive understanding of a species’ history is important for both evolutionary research and for conservation management. However, this may be impossible using data derived solely from living individuals – particularly for endangered species whose current genetic diversity is depauperate. A potential solution is to study DNA sequences obtained from historical or ancient samples [1], allowing extinct populations and lineages to be investigated and compared with those that exist today. In studies that use such samples, mitochondrial DNA continues to be an important marker as the much higher copy number per cell compared to the nuclear genome generally results in a higher success rate for sequence recovery - particularly for species with low fossil abundance and/or poor biomolecular preservation [e.g. 2–5]. The leopard (*Panthera pardus* Linnaeus, 1758) is an example of a species that is currently distributed across only a fraction of its historical and ancient range [e.g. 6–8]. It is one of only a few large-bodied carnivore species that naturally occurs in a wide variety of habitats; from the Himalayan highlands to the Ethiopian desert, and from the Congo rainforests to the Amur taiga [9]. Human persecution and hunting [e.g. 10–12], habitat destruction [e.g. 13–15] and reduced prey availability [16, 17] have severely impacted the distribution of this elusive predator, and leopards are now extinct in large parts of their historic Asian and African distribution (Fig. 1) [7, 18].

**Fig 1:**
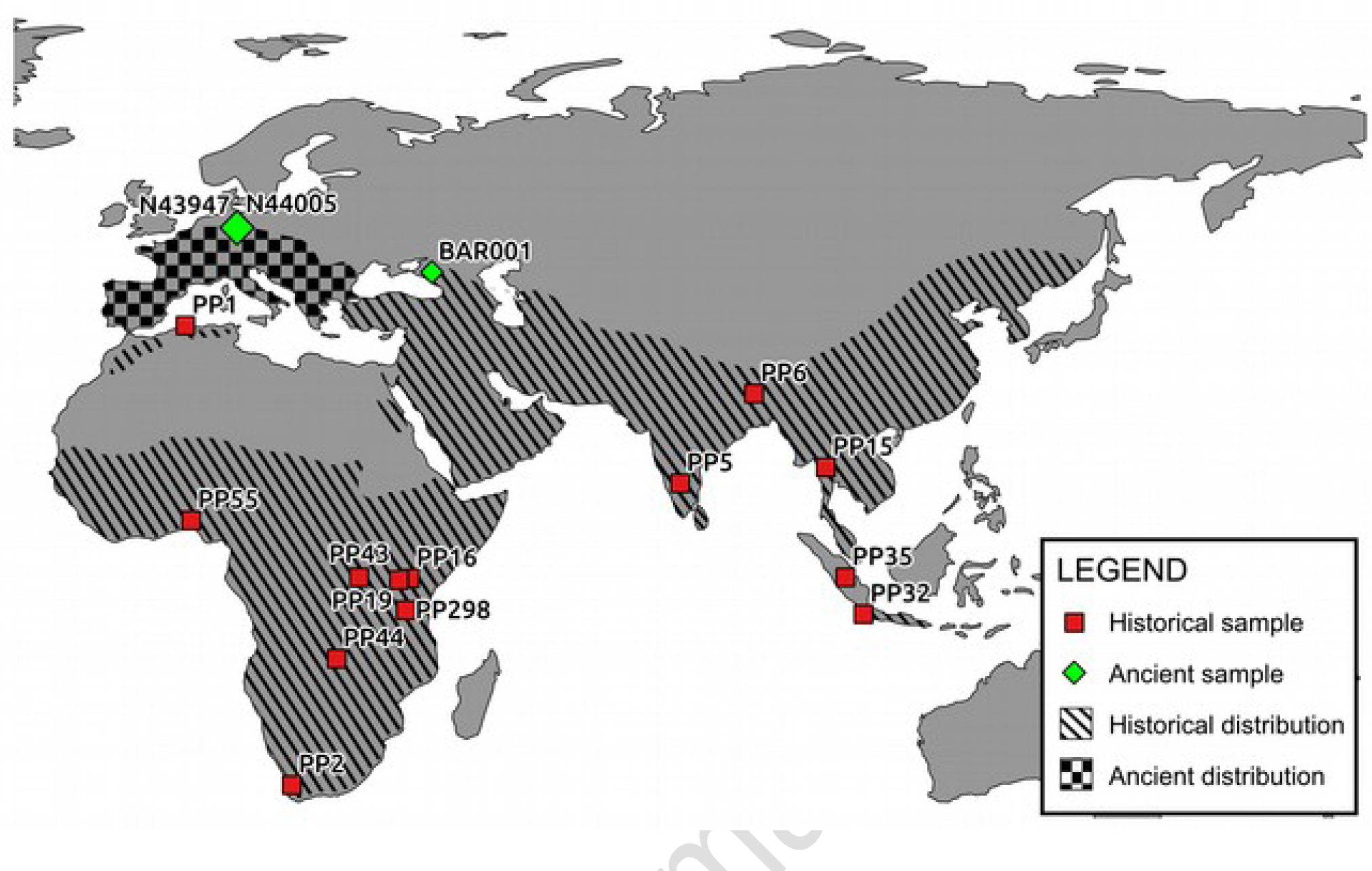
Map indicating the location of the samples included in our study, with the approximate historical and ancient (Pleistocene) distribution of the leopard (adapted from Uphyrkina et al., 2001; Diedrich 2013). The current distribution of leopards is severely reduced compared to the historical range, and highly fragmented (Jacobson et al. 2016). Sample PP3 is not displayed due to its ambiguous provenance (“East Indies”).

The fossil record and genetic data from modern and historical samples are generally interpreted as indicative of an African origin of the leopard. The oldest leopard fossils have been recovered in Eastern Africa, and genetic diversity (estimated using both mitochondrial DNA and nuclear microsatellites) in living African populations is higher than in Asian populations, which has been interpreted as evidence for an African origin [19, 20]. The oldest potential evidence for leopards outside of Africa is found in the Early Pleistocene fossil record from South Asia (Pakistan), suggesting an initial out-of-Africa expansion into Eurasia around 2.0 Ma (*Mega annus*; million years ago; [21]). Mitochondrial and microsatellite data, however, suggest that all current Asian populations share a much more recent common ancestor (approximately 622 Ka [*Kilo annus*; thousand years ago]; [19, 22]); more than a million years younger than the oldest fossils found in Asia. The apparent incongruence between the fossil record in Eurasia (which suggests occupation since the Early Pleistocene) and the relatively recent coalescence time of African and Asian modern leopard mitochondrial lineages has been interpreted as indication for two independent out-of-Africa dispersal events, the latter of which founded all modern Asian leopard lineages [19].

Previous studies that have examined intraspecific variation in leopards have been based on short mitochondrial sequences and microsatellites from recently collected samples [19, 23-29]. In this study, we present near-complete leopard mitogenomes from seven Late Pleistocene specimens (up to 45,000 years old) and from 15 historical samples collected up to 150 years ago, from a range of geographical locations encompassing its entire distribution (Fig. 1). We investigate this mitogenomic data in the context of the proposed evolutionary history and past population dynamics of the leopard, and the role of Pleistocene populations within this scenario. Furthermore, by combining mitochondrial data from previous studies [19, 22, 24, 27] with ancient and historical mitochondrial DNA (mtDNA), we present new insights into ancient and recent population dynamics of one of the most widespread extant felid species.

## Results

Using hybridisation capture and high-throughput sequencing, we retrieved near-complete mitogenomes from 22 leopard samples: one Late Pleistocene Caucasian leopard (∼35 Ka), six Late Pleistocene Central European leopards (∼45 Ka), and 15 historical leopards from across the entire historical distribution (Fig. 1; Table 1). The mitogenome sequences represent 17 distinct haplotypes, which were analysed in conjunction with three previously published modern leopard mitogenomes. Using a two-step approach (see Methods section for details), we estimated the coalescence times of this dataset in BEAST. Our results support the deep bifurcation between Asian and African leopards, that has previously been proposed based on short mtDNA sequences (Fig. 2; Additional File 1: Fig. S1; [19, 22]. The fossil-calibrated Bayesian analysis of the basal divergence time for all leopard mitochondrial lineages was 710 Ka (95% credibility interval [CI]: 457 - 956 Ka), which is also consistent with the previous estimates (932 Ka; Wilting et al 2016, 471 - 825 Ka; [19]). The Pleistocene European sequences form a well-supported clade consisting of three distinct haplotypes, which is sister to a clade containing all mitogenome sequences from Asian leopards. The two clades are estimated to have diverged approximately 483 Ka (95% CI: 305 - 677 Ka). The ancient Caucasian sequence is sister to all modern mainland Asian sequences, with high support (Bayesian posterior probability of 1.0, Fig. 2; Additional File 1: Fig. S1). The estimated coalescence time of mainland Asian mitogenomes, including the ancient Caucasian individual, is 244 Ka (95% CI: 148 - 352 Ka).

**Table 1:** Samples from which a (almost) complete mitogenome was reconstructed or recovered from GenBank. Samples indicated with an asterisk are C14 dated, the date is provided in uncalibrated years before present (additional dating information is included in Additional File 1: Table S3).

| HISTORICAL |  |  |  |  |  |  |
| --- | --- | --- | --- | --- | --- | --- |
| Museum accession number | Sample code | Origin | Date | mtDNA recovery | Collection | GenBank accession number |
| ZMUC 24 | PP1 | Algeria (North Africa) | 1850 | 2x serial capture | Natural History Museum of Denmark | MH588618 |
| ZMUC 25/26/206 | PP2 | South Africa (Cape South Africa) | Unknown | 2x serial capture | Natural History Museum of Denmark | MH588619 |
| ZMUC 27/14 | PP3 | East Indies (South-East Asia) | 1836 | 1x capture | Natural History Museum of Denmark | MH588620 |
| ZMUC 29 | PP5 | India (Central India) | 1839 | Shotgun sequencing | Natural History Museum of Denmark | MH588621 |
| ZMUC 775 | PP6 | India (East India) | 1884 | 1x capture | Natural History Museum of Denmark | MH588622 |
| ZMUC 1967 | PP15 | Thailand (South-East Asia) | 1924 | 1x capture | Natural History Museum of Denmark | MH588623 |
| ZMUC 2115 | PP16 | Kenya (East Africa) | 1928 | 2x serial capture | Natural History Museum of Denmark | MH588624 |
| ZMUC 3490 | PP19 | Kenya (East Africa) | 1945 | Shotgun sequencing | Natural History Museum of Denmark | MH588625 |
| ZMUC 3980 | PP26 | Burundi (Central-East Africa) | 1880 | 1x capture | Natural History Museum of Denmark | MH588626 |
| ZMUC 31 | PP32 | Indonesia (Java) | 1848 | Shotgun sequencing | Natural History Museum of Denmark | MH588627 |
| ZMUC 34 | PP35 | Indonesia (Sumatra) | 1866 | 2x serial capture | Natural History Museum of Denmark | MH588628 |
| ZMUC 3739 | PP43 | DRC (Central Africa) | 1951 | 2x serial capture | Natural History Museum of Denmark | MH588629 |
| ZMUC 4446 | PP44 | Zambia (Central Africa) | 1960 | Shotgun sequencing | Natural History Museum of Denmark | MH588630 |
| ZMUC 5719 | PP55 | Nigeria (West Africa) | 1958 | 1x capture | Natural History Museum of Denmark | MH588631 |
| ZMUC SC007 | PP298 | Tanzania (Eastern Arc) | 2013 | 2x serial capture | Natural History Museum of Denmark | MH588632 |
| ANCIENT |  |  |  |  |  |  |
| Museum accession number | Sample code | Origin | Date | mtDNA recovery | Collection | GenBank accession number |
| - | BAR001 | Mezmaiskaya Cave (Russia, Northern Caucasus) | > 35 Ka | 2x serial capture | Zoological Institute, St. Petersburg | MH588611 |
| N43947 | N43947 | Baumannshöhle (Rübeland, Germany) (Joger & Rosendahl, 2012) | ~ 40 Ka | 2x serial capture | Natural History Museum Braunschweig | MH588612 |
| N43951 | N43951 | Baumannshöhle (Rübeland, Germany) (Joger & Rosendahl, 2012) | ~ 40 Ka | 2x serial capture | Natural History Museum Braunschweig | MH588613 |
| N43962 | N43962 | Baumannshöhle (Rübeland, Germany) (Joger & Rosendahl, 2012) | 44710±630* | 2x serial capture | Natural History Museum Braunschweig | MH588614 |
| N43974 | N43974 | Baumannshöhle (Rübeland, Germany) (Joger & Rosendahl, 2012) | ~ 40 Ka | 2x serial capture | Natural History Museum Braunschweig | MH588615 |
| N43981 | N43981 | Baumannshöhle (Rübeland, Germany) (Joger & Rosendahl, 2012) | 37880±300* | 2x serial capture | Natural History Museum Braunschweig | MH588616 |
| N44005 | N44005 | Baumannshöhle (Rübeland, Germany) (Joger & Rosendahl, 2012) | 40470±410* | 2x serial capture | Natural History Museum Braunschweig | MH588617 |
| PUBLISHED |  |  |  |  |  |  |
| NCBI GenBank accession number | Sample code | Origin |  | Reference |  |  |
| NC_010641 | NC_010641 | China |  | Lei et al. 2011 |  |  |
| KJ866876 | KJ866876 | North-Chinese leopard |  | Dou et al. 2016 |  |  |
| KP202265 | KP202265 | North-Chinese leopard |  | Li et al. 2016 |  |  |

**Fig 2:**
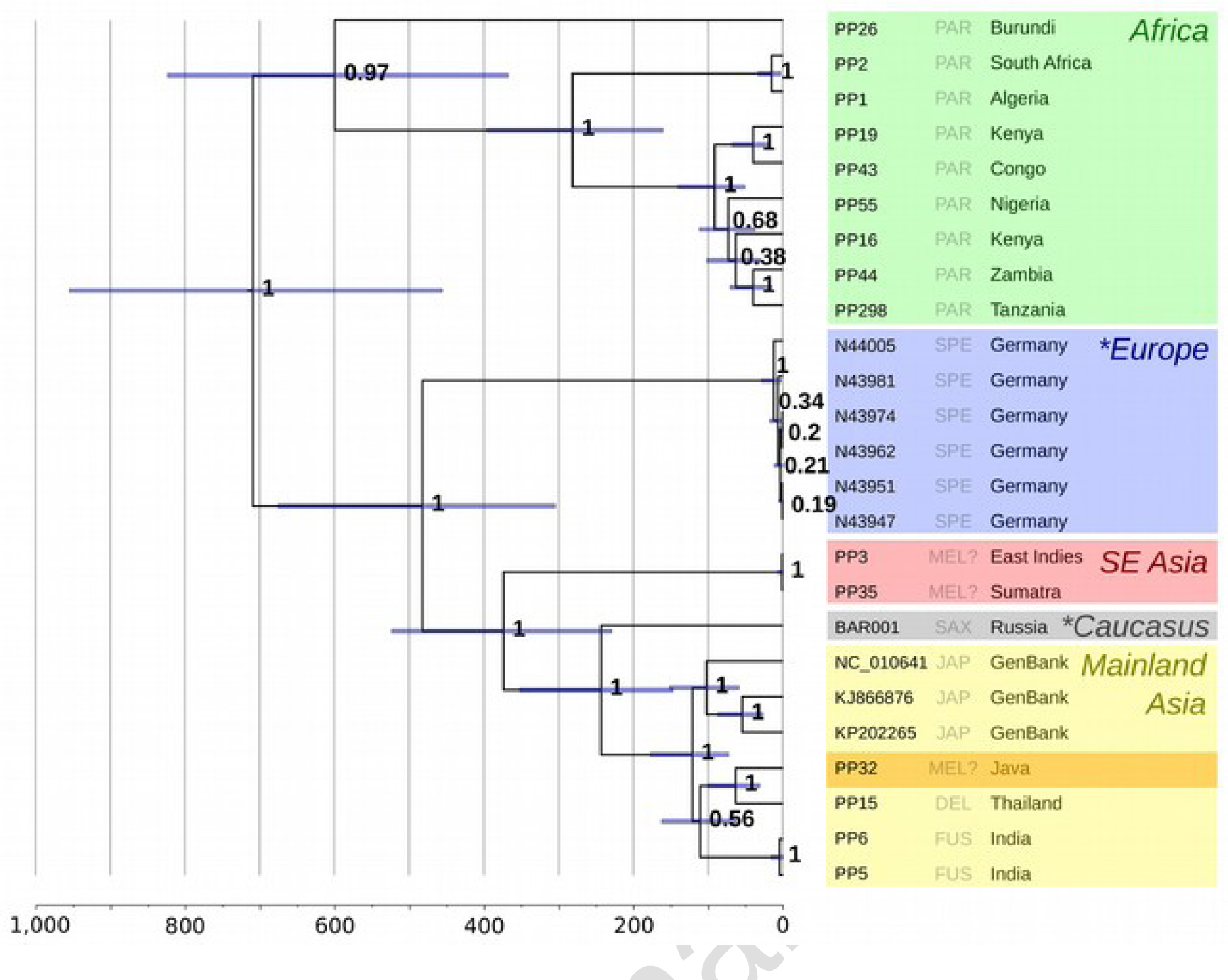
Calibrated mitogenomic phylogeny of 25 leopard mitogenome sequences. Node support is indicated by Bayesian Posterior Probabilities, blue node bars indicate the 95% credibility interval of divergence times. The lower axis shows the estimated coalescence times in thousands of years. Colours indicate the locality of the samples; the unexpected placement of the Javan leopard is highlighted dark yellow. The three-letter code corresponds to the putative subspecies for each individual, following Miththapala *et al.* 1996, Uphyrkina et al., 2001; Diedrich 2013. Asterisks indicate the Late Pleistocene samples. The RaXML maximum likelihood phylogeny can be found in Additional File 1: Fig. S1.

The South-East Asian samples included one specimen collected in Sumatra (PP35), where currently no leopards live and thus is likely to represent a traded specimen imported from elsewhere. A second specimen in this clade had no associated geographical provenance (’East Indies’; PP3). These individuals may represent the Javan leopard lineage, as their mitogenome haplotypes are sister to all Asian leopard sequences (Fig. 2), and they exhibit ND5 sequences identical to previously published sequences from Javan leopards [19, 22]; Additional File 1: Fig. S2). The coalescence time of this lineage and the mainland leopards is estimated at 375 Ka (95% CI: 230 - 524 Ka). In contrast to previous studies, our Javan sample (PP32) was placed as a sister lineage to a specimen from Thailand (PP15), nested within all mainland Asian leopards rather than sister to these, with an estimated coalescence time of 64 Ka (95% CI: 32 - 100 Ka; Fig. 2).

## Discussion

Using modern, historical and ancient mitogenomic data from leopard samples from across their current and former geographic range, we provide novel insights into the historical biogeography of the leopard. The resulting data clarify the relationship between current leopard populations and Late Pleistocene fossils, and provide additional evidence for the interpretation of the earliest putative leopard fossils.

### Origin of the leopard

Based on the fossil record, the origin of the leopard has been placed in Eastern Africa. There are fossils that may belong to *P. pardus* dating to about 3.4 - 3.8 Ma from Laetoli (Olduvai; [30]), although some authors suggest that these may be assigned to a different *Panthera* species [31], or some other large-bodied felid [32]. Although the fossil record between 2.0 and 3.8 Ma is sparse, unequivocally identified leopard fossils do confirm their presence in Eastern Africa around 2 Ma [33], which strongly suggests an African origin of the species (Fig. 3).

**Fig 3:**
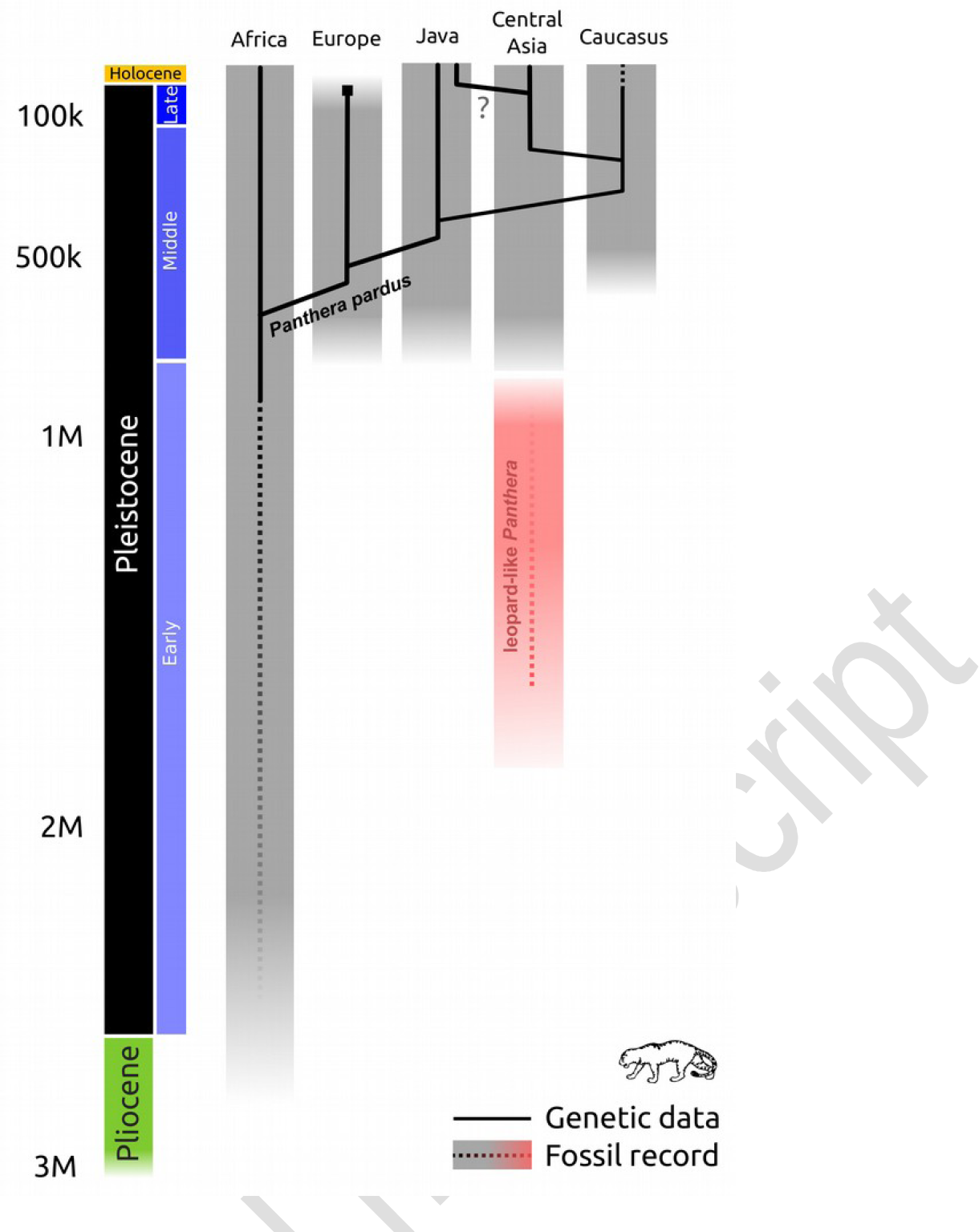
Proposed phylogeographical history for the leopard (*Panthera pardus*), comparing the genetic data with the fossil record. Shaded areas indicate the age of the fossil record for *P. pardus* (grey) for each region, and the disputed Asian leopard-like *Panthera* fossils (red). The solid lines indicate the relationships between the mitogenome lineages recovered in this study.

We estimate the coalescence time between African and Eurasian leopard mitochondrial lineages to be ∼710 Ka, similar to the age found by previous molecular studies [19, 22]. Although the bifurcation between African and Eurasian leopards provides no confirmation for either an African or Eurasian origin for the leopard, the combined fossil and genetic evidence together support Africa as the most likely place of origin. The relatively recent coalescence time between African and Eurasian mitochondrial lineages is also consistent with previous suggestions that the oldest African fossils of 3.8 Ma may not in fact represent *P. pardus* [31, 32]. Within Africa, we found considerable genetic divergence between haplotypes occurring in East Africa. In particular, the haplotype sampled from the Burundi individual forms a divergent sister lineage to all other sampled African haplotypes (95% CI: 368 - 814 Ka), including those from other regions of East Africa. Thus, the mitochondrial divergence occurring within this region of Africa is equivalent to that of the entire African continent, indicating this region as the potential point of origin of the leopard, which has also been suggested based on short mtDNA sequences [23]. Given that, based on the fossil record, leopards were present in East Africa at least around 2 Ma ago [31], this hypothesis requires that mitochondrial lineages established prior to ∼600 Ka have either being lost from modern populations through genetic drift or have not yet been sampled. These results do suggest, however, that, as well as for the genus *Homo* and a number of other species [34], East Africa may be the point of origin of modern leopards, although additional sampling of individuals and of nuclear markers is desirable to more robustly test this hypothesis.

### Out-of-Africa dispersal

The first evidence for leopards outside of Africa is ambiguous. In Asia, the earliest occurrence of leopard fossil remains is in the Early Pleistocene of South Asia (Pakistan), suggesting an initial out-of-Africa expansion into Asia around 2.0 Ma [21]. Whether or not these fossils should be assigned to *P. pardus* is subject to discussion, however [31, 32]. These ancient specimens from Pakistan may be better assigned to other medium-sized felids (e.g. Eurasian puma, jaguar or snow leopard), or indicate an earlier dispersal of a leopard-like *Panthera* taxon into Asia (Fig. 3). Unequivocal leopard fossils in Asia are much younger; 0.6 – 0.8 Ma [35, 36]. Furthermore, in Europe, the oldest findings of this species are dated to the early Middle Pleistocene (nearly 0.6 Ma: [37–39].

Inferring the timing of colonisation events from the mitochondrial gene tree is challenging, as the population divergence and lineage coalescence will be asynchronous. Lineage coalescence of founding and source populations represents the maximum limit for dispersal time, but could be an overestimate caused by deep coalescence within the source population. Assuming monophyly of founding population lineages, the radiation of the founding population represents the minimum limit on the dispersal time, but this will tend to be an underestimation due to the mutation-lag of the formation of new lineages, incomplete sampling of extant lineages, or by lineage extinction during population bottlenecks. Thus, the true colonisation event lies somewhere along the branch connecting these upper and lower limits. Following this reasoning, the mitogenome data suggest the colonisation of Eurasia (or at least the end of maternal gene flow) some time between 710 and 483 Ka ago. This range overlaps with the age of the younger, unequivocal leopard fossils from Asia but not with the older Early Pleistocene Asian fossils whose precise taxonomic assignment has been debated. Achieving a unified biogeographic hypothesis for the colonisation of Asia by leopards thus hinges on the classification of these older fossils. Under the assumption that the first leopard occurrence in Eurasia is represented by the unequivocally identified specimens dating to around 0.6 - 0.8 Ma, there is no contradiction between the fossil record and the molecular evidence. If, however, older Early Pleistocene specimens are to be classified as *P. pardus*, then multiple out-of-Africa dispersals and the replacement of all mitochondrial lineages present in Asia from earlier dispersal events is required to explain the observed patterns. Considering the ambiguity of the Early Pleistocene fossil record in Asia, we consider a single out-of-Africa for *P. pardus* more parsimonious given the currently available combined palaeontological and genetic evidence. Clarifying whether the African and Eurasian leopard populations have indeed been separated since this time, or if gene flow continued to some extent, and if the earlier Eurasian leopard-like populations played a role in the evolutionary history of the more recent populations, will require nuclear genomic data with a similar geographical coverage.

### Ancient Eurasian leopards

The origin and classification of the European Pleistocene leopard is equivocal based on fossil remains from the region [e.g. 6, 40–42]. The fossil record suggests that leopards first dispersed into Europe during the Middle Pleistocene, as evidenced by fossils from Western Europe estimated to be around 600 Ka [6, 30, e.g. 35, 37, 38]. Several different forms have been defined in European fossil assemblages based on morphology, which has been interpreted as evidence for several immigration waves from Africa or Central Asia [6]. The Late Pleistocene form, representing the age of the specimens we retrieved mitogenomes from, has been referred to as the subspecies *P. p. spelaea [6, 42].* These populations likely went extinct before the Last Glacial Maximum (LGM), around 25,000 years ago [6, 8, 40, 41, 43]. The disappearance of the European leopard populations coincides with the extinction of many other Pleistocene carnivores, such as the cave bear (*Ursus spelaeus*), cave hyena (*Crocuta crocuta spelaea*) and *Homotherium* [3, 44-46].

We recover three distinct haplotypes from the six leopard samples, and as there are no overlapping skeletal elements in the sample set, the number of individuals sampled is uncertain (minimum of three). We find that the Late Pleistocene European leopards share a more recent common ancestor with modern Asian leopards, than either does with African leopards (Fig. 2; Additional File 1: Fig. S1). The estimated coalescence time between the European and Asian mitogenomes is 485 Ka, with a 95% CI of 301 - 675 Ka, which encompasses the age for the oldest fossil records of Europe. This suggests that the European leopards originate from the same out-of-Africa event as the Asian leopard lineages, rather than from an independent migration. The ancient Caucasian leopard lineage is sister to all modern samples from mainland Asia, with an estimated coalescence time of ∼244 Ka (95% CI: 148 - 352 Ka).

Morphological studies have failed to differentiate Pleistocene Eurasian leopard fossils from their modern counterparts [21]. Cranial and postcranial measurements from Middle Pleistocene remains from Poland (estimated to 569 - 125 Ka; [39] and from the Caucasus (less than 350 Ka; [37]) overlapped with the variation found in modern leopards, which has been interpreted as evidence for some degree of population continuity between those ancient and modern populations. The dates of these fossils are not older, and actually correspond fairly well to the estimated coalescence time between ancient European mitochondrial lineages and modern leopards (95% CI: 305 - 677 Ka) or between the ancient Caucasian individual and modern leopards (95% CI: 148 - 352 Ka). Thus, some degree of population continuity since the Late Pleistocene, as suggested by the morphology, is not in conflict with our molecular dating. Alleles of Pleistocene European leopards may thus survive in the modern Caucasian population. However, as our dataset does not include modern Caucasian individuals, nor ancient East Asian ones, a direct assessment of population continuity between the Middle Pleistocene and today can only be based on short mtDNA sequences [24, 27]; Additional File 1: Fig. S2). The mtDNA haplotype network has only limited resolution as it is only based on 456 bp of mtDNA, but does not support that the ancient Caucasus leopard is particularly diverged from modern Persian leopards (*P. p. saxicolor*; “SAX”; Additional File 1: Fig. S2). The discrepancy between the coalescent times reported for the divergence between Iranian and other Asian leopards (16 – 270 Ka; [24]), and that between our Caucasian sample and other lineages (148 – 352 Ka) could suggest that the ancient Caucasian leopard represents a distinct lineage that diverged from other Asian leopards prior to the Iranian leopards, although it should be noted that the credibility intervals overlap considerably. Tracing the potential contribution of the European leopard to modern Caucasian or other modern leopard populations will require the analysis of nuclear DNA from Pleistocene leopards. Unfortunately, the Pleistocene samples analysed in the present study were not suitable for the analysis of nuclear DNA due to their low endogenous DNA content, despite multiple re-sampling attempts [47].

### Java and the Asian leopard

The Javan leopard has been assigned to the subspecies *P. p. melas* (“MEL”), based on both morphological and genetic distinctiveness of these island leopards from mainland Asian populations [19, 22, 36]. Previous studies found Javan leopards to represent a mitochondrial lineage that is sister to the entire diversity of mainland Asian leopards [19, 22]. Due to this position as sister lineage to all Asian leopards, it has been suggested that the Javan leopard represents a relict population founded from a Middle Pleistocene leopard population from the South East Asian mainland, after which it became isolated on the Sundaic islands (Uphyrkina et al., 2001; Wilting et al., 2016). Deep divergence between mainland and Sundaic species or populations has also been found in a range of other animals including terrestrial mammals [48] such as tigers [49] and leopard cats [50], but also birds (Hughes et al., 2003), bats (Hughes et al., 2011), and frogs (Inger and Voris, 2001).

Under the assumption that both of our samples with unclear provenance (‘Sumatra’ and ‘South East Asia’, sample codes PP3 and PP35) do in fact represent the Javan population, the molecular dating performed with our mitogenome dataset suggests that the Javan leopards became established on the island some time after 375 Ka (95% CI: 230 - 524 Ka). This is in line with previous estimates (393 - 886 Ka; [22]). Low sea levels during glacial periods exposed portions of the Sunda shelf, creating land bridges connecting the mainland with the Sundaic islands [51], which would have allowed the colonisation of and subsequent isolation on the South East Asian islands [22, 36]. If there was any more recent gene flow between the islands and the mainland during Late Pleistocene periods of low sea levels [52], it is not expected to have been at high frequency, as there has been no evidence found in the mitochondrial data investigated until now. However, the sample in our dataset that was reported to be of Javan origin (sample PP32) was placed as a sister lineage to the mitogenome recovered from an individual from Thailand. Investigation of short mtDNA sequences confirmed that the mitochondrial haplotype from this putative Javan leopard is closely related to *P. p. delacouri* (“DEL”), the mainland South-East Asian leopard subspecies (Additional File 1: Fig. S2). Trade routes between the South East Asian islands and the mainland have been reported to exist for almost 2000 years [53], so leopards could have been moved to Java via such human-mitigated routes – either by trading live animals or as hunting trophies. Alternatively, our results also fit a scenario in which leopards migrated to the South-East Asian islands from the mainland during the late Pleistocene, supplementing the original locally surviving Javan mitochondrial haplotypes with mainland Asian haplotypes; the latter of which could have been lost during the drastic population declines during the past century [7]. The coalescence time of this putative Javan leopard and the Thai individual is estimated to be 64 Ka (95% CI: 32 - 100 Ka), which could be consistent with a dispersal from the mainland to the Sundaic islands some time (e.g. during the LGM) after the Toba supervolcanic eruption that occurred 73 Ka [54]. Additional sampling and/or the investigation of nuclear data is needed to further clarify the origin and relatedness between mainland and Javan leopards.

Our dataset further included a number of South and East Asian leopards; two Indian leopards (*P. p. fusca*), one Indochinese *P. p. delacouri* and three published North-Chinese leopards, *P. p. japonensis* (Table 1; “FUS”, “DEL” and “JAP”, respectively). The estimated coalescence time for these individuals is relatively recent (122 Ka; 95% CI: 73 - 178 Ka). This recent divergence time could point towards extensive population reductions on the Eurasian mainland during the Pleistocene, leading to a bottleneck and loss of divergent lineages for all Eurasian leopards except for Java and Europe. A loss of genetic diversity during the Pleistocene as suggested by recent mitogenome coalescence times has also been found for other carnivores (e.g. 151 Ka for leopard cats, 95% CI: 87 - 215; [50]), suggesting that this pattern may be detectable across other Asian carnivores. Considering the deeper coalescence time of all modern African mtDNA lineages (600 Ka), the African leopards are not likely to have experienced a bottleneck to the same extent as Asian leopards.

## Conclusions

This study represents an example of how genetic data from ancient DNA can inform on the evolutionary history of species and thereby provide additional clues on identification of fossils, including those from which no genetic data can be recovered because they are beyond the expected range of DNA survival, such as the 2 Ma year old fossils from Pakistan. The molecular dating of mitochondrial lineages is in line with the age of the earliest unequivocal leopard fossils from Eurasia, and thus it encourages a re-evaluation of the older leopard-like fossils. Our results are therefore compatible with previous suggestions that older fossils should be assigned to other large-bodied felids. Furthermore, similar to many other Eurasian carnivores, our data suggests that with the extinction of European populations and the proposed Pleistocene population bottleneck in mainland Asia, leopards experienced not only a reduction of their geographical distribution but also a loss of mitochondrial lineages that, to date, have not been detected in the modern gene pool. Leopard dispersal is generally driven by males, and thus may not be detectable using mitochondrial DNA alone. Finally, although we do not observe any evidence in our data to suggest that ancient European mitochondrial lineages persist in modern Asian populations, it is possible that at least part of the genetic legacy of Pleistocene leopards survives in the modern nuclear genome, as was recently shown to be the case for archaic hominins [55, 56] and the extinct cave bear [57].

## Methods

### DNA extraction, library preparation and capture

All pre-PCR steps for ancient and historical bone samples were performed in dedicated cleanroom facilities of the Evolutionary Adaptive Genomics group at Potsdam University, with all standard ancient DNA precautions (e.g. decontamination procedures for all reagents and materials, negative controls for both extraction and library preparation). Extraction was performed following a protocol optimised for the retrieval of short DNA fragments [58]. Illumina sequencing libraries were constructed following a double-stranded library preparation protocol for the historical samples [59, 60] and a single-stranded library protocol for the ancient samples, using UDG/Endonuclease VIII to excise uracils from the molecules [61]. Optimal amplification cycle numbers for each library were estimated using qPCR and then used for dual-indexing PCR. Four historical samples yielded high endogenous content, allowing the mitogenome sequence to be recovered using shotgun sequencing (Table 1; Additional File 1: Table S1). For all remaining samples, mitogenome enrichment was performed using an in-solution capture approach with synthetic baits. A bait-set targeting felid mitochondrial DNA was designed by placing 52-mer probes (1 bp tiling) across the mitogenomes of five felids: leopard (*Panthera pardus*, EF551002 (RefSeq: NC_010641), lion (*Panthera leo*, KF776494), domestic cat (*Felis catus*, NC_001700), bobcat (*Lynx rufus*, NC_014456.1) and cheetah (*Acinonyx jubatus*, NC_005212). Bait sequences containing simple repeats longer than 24 bp were removed [62]. Baits were synthesized, amplified and converted into biotinylated single-stranded DNA probes as described elsewhere [63]. Capture was performed following the protocol in Horn (2012). For ancient samples, two serial captures were performed to improve the enrichment rates. For historical samples, either one or two captures were used (Table 1). Sequencing was performed on the Illumina NextSeq platform using 75bp paired-end sequencing, using custom sequencing primers for the single-stranded libraries [61, 64].

### Sequence processing

Raw sequences were trimmed and merged using SeqPrep with default parameters and a minimum length of 30bp (available from https://github.com/jstjohn/SeqPrep). The trimmed and merged reads were then aligned to a leopard mitogenome sequence available from GenBank (Acc. Nr. KP202265) using the Burrows-Wheeler Aligner (BWA aln) v0.7.8 and samtools v1.19 [65, 66] with default parameters. Reads with low mapping quality (<Q30) were removed. Duplications were marked and removed taking both mapping coordinates into consideration, using MarkDupsByStartEnd.jar (http://github.com/dariober/Java-cafe/tree/master/MarkDupsByStartEnd). Summary statistics can be found in Additional File 1: Table S1. Mitogenome consensus sequences were retrieved using Geneious v7.0 [67], using a minimum sequence depth of 3x and a strict 90% majority rule for base calling. The resulting consensus sequences for each sample were combined with three mitogenome sequences available from GenBank at the time of analysis, and aligned using ClustalW with default parameters ([68]; as implemented in Geneious). The control region, as well as any positions in the alignment that contained missing or ambiguous data, were removed to avoid any adverse influence of misalignments or numts. The resulting alignment (13,688 bp in length) was manually annotated in Geneious using the published sequence (*Panthera pardus*; Genbank Acc. Nr. KP202265) as reference.

### Phylogenetic analyses

PartitionFinder v1.1.1 [69] was used to identify an optimal partitioning scheme from all possible combinations of tRNAs, ribosomal RNA genes and protein-coding genes, considering all substitution models available in BEAUti, using the Bayesian Information Criterion (BIC). The partitioned maximum likelihood tree was calculated using RaxML-HPC v8.2.4 [70] on the CIPRES black box version, with default substitution models for each partition, on the CIPRES Science Gateway ([71]; Additional File 1: Fig. S1). Molecular dating was performed using BEAST v1.8.2 [72]. We adopted a two step strategy for the calibrated analyses: first, we generated a fossil-calibrated phylogeny using a total of seven fossil dates (Additional File 1: Table S2) under a Yule speciation model for 15 sequences from various Felidae species, including two of the most divergent leopard mitogenomes. The divergence estimates recovered for the two leopard mitogenomes (mean 0.81 Ma, standard deviation 0.12 Ma) was then applied as normal prior on the root height for the intraspecific analyses, and were run with a Bayesian Skyline coalescent population model. Preliminary analysis using a lognormal relaxed clock model failed to reject zero variation in substitution rates across branches of the phylogeny, and so a strict clock model was employed with an open uniform prior on the mean per-lineage substitution rate of 0 to 20% per million years. The MCMC chain was run for a sufficient number of generations to achieve convergence (to a maximum of 10 million, sampling every 10,000 states) and adequate posterior sampling of all parameters (ESS >200), checked using Tracer v1.5 (available from http://www.beast.bio.ed.ac.uk/Tracer). TreeAnnotator v1.8.2 was then used to remove the first 25% of trees as burnin (corresponding to 2,500 trees) and extract the Maximum Clade Credibility (MCC) tree with nodes scaled to the median heights recovered by the posterior sample. The minimum-spanning network for the short mtDNA alignment (267 individuals, 456 bp in length) was generated using Popart (Additional File 1: Fig. S2) [73].

## List of abbreviations

Ma –: *Mega annus*; million years ago
Ka –: *Kilo annus*; thousand years ago
mtDNA –: mitochondrial DNA
CI –: Credibility Interval
LGM –: Last Glacial Maximum

## Declarations

### Ethics approval and consent to participate

Not applicable

### Consent for publication

Not applicable

### Availability of data and material

Leopard mitogenome consensus sequences will be made available on GenBank (NCBI Accession Numbers MH588611-MH588632). The alignment used for phylogenetic inference and the BEAST XML input file used for fossil calibration analyses will be made available upon request.

## Authors’ contributions

JLAP, MH conceived the project; JLAP, MM designed the experiments; JLAP, KH, BN performed the experiments; JLAP, AB analysed the data; GFB, RWH, DN, UJ, WR, contributed samples and provided interpretation of their context, JLAP, MH, AB, DWF, GFB participated in fundamental discussion and interpretation of the results; JLAP and AB wrote the paper with input from all authors. All authors have read and approved the manuscript.

## Acknowledgements

We would like to acknowledge Peter Rask Møller and Hans Baagøe from the Natural History Museum of Denmark for providing access to samples from their zoological collection. We thank Mathias Stiller for help with the capture bait design. We also acknowledge Tom Gilbert for supervision of RWH, and for providing helpful comments on earlier drafts of the manuscript.

## Additional File 1

**Fig. S1:** Maximum likelihood phylogeny, with different *Panthera* species as outgroup.

**Fig. S2:** Minimum-spanning network of short (456 bp) mtDNA sequences (ND5 gene), including previously published data (based on 267 sequences in total). Number of substitutions are indicated as tick-marks on the branches connecting the haplotypes. Colours indicate the subspecies [19, following 25].

**Table S1**: Summarised sequence statistics for samples included in our study.

**Table S2**: Fossil constraints and calibration priors used in the time-calibrated BEAST analysis performed for Felidae alignment. The resulting root age was then applied as calibration for the leopards-only phylogeny.

**Table S3:** Radiocarbon dating (^14^C) information.

